# Cryo-EM structure revealed a novel F-actin binding motif in a *Legionella pneumophila* lysine fatty-acyltransferase

**DOI:** 10.1101/2025.04.18.649563

**Authors:** Wenjie W. Zeng, Garrison Komaniecki, Jiaze Liu, Hening Lin, Yuxin Mao

## Abstract

*Legionella pneumophila* is an opportunistic bacterial pathogen that causes Legionnaires’ disease. To establish an intracellular niche conducive to replication, *L. pneumophila* translocates a diverse array of effector proteins that manipulate various host cellular processes, including the actin cytoskeleton. In a screen for effectors that alter actin dynamics, we identified a *Legionella* effector, Lfat1 (lpg1387), which colocalizes with the actin cytoskeleton in eukaryotic cells. Lfat1 specifically binds F-actin through a novel actin-binding domain (ABD). High-resolution cryo-electron microscopy (cryoEM) analysis revealed that this ABD forms a long α-helix hairpin, with its tip interacting with subdomains I and II of two adjacent actin molecules within the F-actin filament. Interestingly, while individual α-helices of the hairpin fail to bind F-actin, co-expression as separate fusion proteins restores binding activity. Furthermore, we demonstrated that Lfat1 exhibits lysine fatty-acyltransferase (KFAT) activity, targeting host small GTPases. These findings establish a foundation for studying the KFAT family of bacterial toxins and uncover a novel F-actin binding motif, providing an alternative F-actin marker with notable flexibility.

## Introduction

The actin cytoskeleton plays an essential role in diverse cellular processes, including cell motility, cytokinesis, intracellular trafficking, and cell signaling (Dominguez & Holmes, 2011; Letort *et al*, 2015; Pollard, 2016). Actin is one of the most conserved, ubiquitous, and abundant proteins in cells from amoebas to humans (Pollard, 2016). Actin exists in two distinct forms: the monomeric G-actin form and the double-stranded filamentous F-actin form. F-actin is highly dynamic with a net association of ATP-actin to the barbed (+) end and dissociation of ADP-actin monomers from the pointed (-) end (Pollard, 2016). The assembly and disassembly of F-actin *in vivo* is intricately regulated through interactions with a structurally and functionally diverse family of Actin-Binding Proteins (ABPs) (Pollard, 2016). These actin-binding proteins have usually been classified according to their functional properties into several families, such as the actin monomer-binding protein profilin (Carlsson *et al*, 1977); actin nucleators, including Arp2/3 and Formin that initiate *de novo* branched and unbranched filament assembly, respectively (Chen *et al*, 2010; Dong *et al*, 2003; Machesky *et al*, 1994; Pizarro-Cerda *et al*, 2017); the heterodimeric capping proteins that terminate F-actin elongation (Isenberg *et al*, 1980); severing/depolymerization factors Cofilin and Gelsolin (Bamburg *et al*, 1980; Barrie *et al*, 2024; Tanaka *et al*, 2018; Yin & Stossel, 1979); filament binding proteins, such as tropomyosin that binds to the main groove of an actin filament and stabilizes the filament (von der Ecken *et al*, 2015; Yu & Ono, 2006); and cross-linking proteins that crosslink and stabilize multiple F-actin filaments together for cell movement and muscle contraction (Le *et al*, 2017; Ribeiro Ede *et al*, 2014). The accumulation of 3-dimensional structures of ABPs in complex with actin revealed that different ABPs share a limited number of actin-binding structural modules (Van Troys *et al*, 1999). Thus, identifying and characterizing new actin-binding structural modules will provide direct hints of the actin target site and the functional effect on the actin dynamics of ABPs that share the specific actin-binding module.

Given actin’s essential roles in eukaryotes, many prokaryotic and viral pathogens co-opt a variety of mechanisms that target the host actin cytoskeleton for effective pathogenesis (Aktories *et al*, 2011; Bugalhao *et al*, 2015). The virulence factor BimA from *Burkholderia pseudomallei* mimics host actin-polymerizing proteins Ena/VASP to nucleate, elongate, and bundle filaments (Benanti *et al*, 2015). The *Vibrio parahaemolyticus* VopL consists of a VopL C-terminal domain (VCD) and three WASP homology 2 (WH2) motifs and mimics the Arp2/3 complex and formin proteins to stimulate actin polymerization (Namgoong *et al*, 2011; Zahm *et al*, 2013). The *Salmonella* invasion protein A (SipA) effector is an actin-binding protein that enhances actin polymerization and promotes the uptake efficiency of the bacterium (McGhie *et al*, 2004).

The Gram-negative bacterium *Legionella pneumophila* is the causative agent of a potentially fatal form of pneumonia in humans named Legionnaires’ disease (Cunha *et al*, 2016; Fraser *et al*, 1977; McDade *et al*, 1977; Mondino *et al*, 2020). Upon entry into human alveolar macrophage cells, the facultative intracellular pathogen translocates more than 350 different bacterial proteins, known as effectors (Burstein *et al*, 2009; Huang *et al*, 2011; Zhu *et al*, 2011). These effector proteins subvert multiple conserved eukaryotic pathways, such as ubiquitination (Price & Abu Kwaik, 2021; Tomaskovic *et al*, 2022), autophagy (Choy *et al*, 2012; Omotade & Roy, 2020; Thomas *et al*, 2020; Wan *et al*, 2024), lipid metabolism (Hsu *et al*, 2012; Swart & Hilbi, 2020; Toulabi *et al*, 2013), and the actin cytoskeleton (Franco *et al*, 2012; Prashar *et al*, 2018; Zhang *et al*, 2023) to aid the pathogen in establishing a *Legionella*-containing vacuole (LCV) amenable to intracellular growth and proliferation (Gomez-Valero *et al*, 2019; Mondino *et al*., 2020; Oliva *et al*, 2018).

Like other bacterial pathogens, *L. pneumophila* utilizes a cohort of virulent effectors to modulate actin. Recent studies revealed that the VipA effector nucleates actin and disrupts the Multi-Vesicular Bodies (MVB) pathway (Franco *et al*., 2012) and the RavK effector cleaves actin to abolish actin polymerization (Liu *et al*, 2017). In our recent screen for *L. pneumophila* effectors that affect host F-actin dynamics, we identified several novel effector proteins that exhibited various degrees of F-actin-associated phenotypes. Among these positive hits, Lpg1387, an effector with no known function, showed strong colocalization with F-actin. In this study, we report the identification of a novel actin-binding motif consisting of a long anti-parallel α-helical hairpin. We further revealed the molecular mechanism of actin-binding by cryo-electron microscopy (cryoEM) and presented evidence for developing a potential F-actin probe based on this novel actin-binding motif. Moreover, using click chemistry, we confirmed that, in addition to the actin-binding motif, Lpg1387 has a lysine fatty-acylate (KFA) catalytic domain specific for small GTPases. Hence, we named this *L. pneumophila* effector Lfat1 (*<u>L</u>egionella F-actin-binding fatty-<u>a</u>cyl-transferase <u>1</u>*).

## Results

### The *Legionella* effector Lfat1 directly interacts with F-actin via a coiled-coil domain

To explore how the intracellular bacterial pathogen *Legionella pneumophila* modulates host actin dynamics, we performed a screen to search for effectors that perturb host actin structures. In this screen, we imaged F-actin structures with phalloidin staining in HeLa cells transfected with an GFP-effectors library. In this screen, several effectors showed various degrees of F-actin-associated phenotypes (Figure S1). Among the positive hits, MavH (Lpg2425) has recently been shown to polymerize actin filaments in a membrane-dependent manner (Zhang *et al*., 2023). Another effector, Lfat1 (Lpg1387), exhibited nearly a complete colocalization with F-actin (Figure S1).

To elucidate the molecular mechanism of how Lfat1 localizes to F-actin filaments, we first analyzed the 3-D structure predicted by AlphaFold (Jumper *et al*, 2021). The structure revealed a hammer-like structure for the full-length Lfat1 protein (Figure 1A). The head of the hammer is formed by a globular NC domain (N and C-terminal globular domain), which is contributed by both the N-terminal (residue 1-137, red) and the C-terminal lobes (residues 356-469 pink) of the protein, while the handle of the hammer is formed by an elongated, anti-parallel, coiled-coil hairpin (CC-domain), which contains the middle portion of the protein (residues 138-355, cyan). To map the region responsible for Lfat1 F-actin localization, we created constructs expressing the NC- and CC-domains fused with an N-terminal GFP, respectively, and investigated their intracellular localization by fluorescence microscopy. Interestingly, while the NC-domain showed a diffused cytosolic localization, the CC-domain exhibited a high colocalization to actin filaments comparable to the wild-type protein (Figure 1B and C). This result was further confirmed by an immunoprecipitation (IP) experiment wherein full-length Lfat1 and the CC-domain were able to pulldown actin whereas the NC-domain could not (Figure 1D). To test whether the CC-domain directly binds actin, we performed an *in vitro* F-actin co-sedimentation assay, in which purified recombinant proteins of the CC-domain were incubated with actin in the presence of G-actin or F-actin buffer. Following ultracentrifugation to pellet F-actin, the supernatant and pellet were analyzed on SDS-PAGE (Figure 1E). In the G-actin buffer, both actin and the CC-domain protein remained in the supernatant; however, the CC-domain protein co-sedimented with the polymerized F-actin in the F-actin buffer, indicating that the CC-domain of Lfat1 directly binds to F-actin. Strikingly, when the CC-domain protein was incubated with actin at a 1:1, 1:2, or 1:4 (actin : CC) molar ratio in the F-actin buffer, an approximately equal amount of CC-domain proteins were co-sedimented with F-actin, and the excess CC-domain proteins remained in the supernatant. This observation indicates that the interaction between the CC-domain and actin is saturable, and the binding occurs at a one-to-one molar ratio (Figure 1F). Together, our findings identified Lfat1 as a novel actin-binding effector of *Legionella*. We further demonstrated that Lfat1 binds F-actin at a one-to-one stoichiometry through a unique, long coiled-coil hairpin CC-domain, which we will henceforth call the actin-binding domain (ABD). .

**Figure 1.**
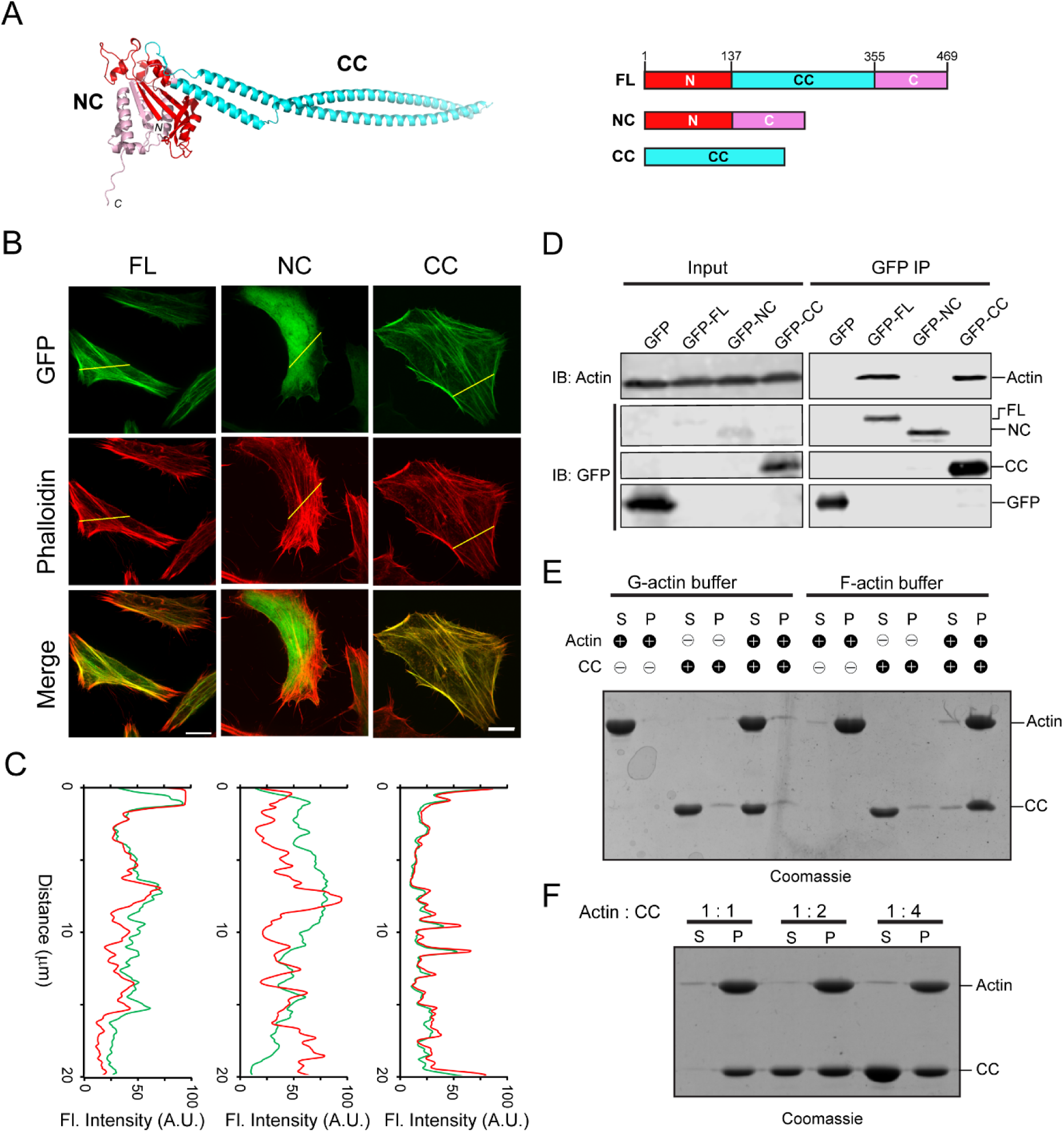
Identification of LFat1 (lpg1387) as an F-actin binding effector. (A) AlphaFold-prediction structure of Lfat1 (Left) and domain architecture of Lfat1 (Right) with the N-terminal domain shown in red, the C-terminal domain in pink, and the central coiled-coil domain in cyan. FL: Full-length, NC: N and C globular domain, CC: Coiled-coil domain, respectively. (B) Cellular localization of Lfat1-FL, -NC, or -CC as determined by fluorescence microscopy. HeLa cells transiently expressing GFP-fused Lfat1-FL, -NC, or -CC were fixed and stained with phalloidin. Scale bar = 10 µm. (C) Colocalization was determined by fluorescence intensity line scan along the yellow line shown in (B). Red = F-actin, Green = GFP. (D) Co-immunoprecipitation to determine interaction of Lfat1 with actin. HEK293T cells transiently expressing either GFP-empty vector, -Lfat1 FL, -Lfat1 NC, or -Lfat1 CC were lysed, and cell lysates were immunoprecipitated using anti-GFP nanobeads. The IP samples were analyzed with SDS-PAGE followed by immunoblot against GFP and actin. (E) Co-sedimentation assay to determine direct interaction between Lfat1 CC with F-actin. Purified G-actin, CC, or G-actin plus CC was incubated either in G-actin buffer or F-actin polymerization buffer, then ultra-centrifuged to separate supernatant from pellet, followed by analysis via SDS-PAGE. S: supernatant, P: pellet. (F) Binding stoichiometry between Lfat1 CC and actin as determined by co-sedimentation assay.

### Cryo-EM structure of the Lfat1 ABD-F-actin complex

The discovery of a novel ABD triggered us to interrogate the molecular mechanism of actin-binding by this ABD. We sought to determine the cryo-electron microscopy (cryo-EM) structure of the Lfat1 ABD-F-actin complex. In our initial attempts to prepare the protein complex, F-actin bundles were readily induced by Lfat1 ABD with the F-actin buffer, making it hard to solve a single F-actin filament for structural determination (data not shown). To restrict excessive actin polymerization, equal molar of Lfat1 ABD and G-actin were incubated in a non-polymerizing G-actin buffer overnight at 4°C. The protein complex sample was then applied to cryo-grids, vitrified, and loaded to a 200kV Thermo Fisher Talos Arctica transmission electron microscope for Cryo-EM data collection. The data were processed, and a high-resolution density map (average to 3.5 angstroms resolution) was calculated and refined using CryoSPARC (Punjani *et al*, 2017). The atomic model of the complex was built by docking the F-actin structure (PDB: 7BTI) and the AlphaFold-predicted model of Lfat1 ABD into the CryoEM density using ChimeraX (Meng *et al*, 2023). The model was then refined iteratively using Phenix (Liebschner *et al*, 2019), and the final model was validated online by wwPDB validation server at https://validate.wwpdb.org (Figure S2 and Supplemental Table 1).

The CryoEM density map of the complex allowed a complete resolution of the actin subunit in the F-actin filament, including its bound ADP and Mg^2+^ ion (Figure 2A, Figure S3A and B). The D-loop (DNase I-binding loop) of the actin monomer adopts a closed conformation. The D-loop of the n^th^ actin monomer extends into the hydrophobic cleft between actin subdomains 1 and 3 of the n+2^nd^ actin monomer. The hydrophobic residues (V45, M46, V47, and M49) at the tip of the D-loop pack against a large hydrophobic area lining the wall of the hydrophobic cleft of the n+2^nd^ actin monomer (Figure 2B). The D-loop also mediates specific hydrogen bond interactions between the two adjacent actin monomers. The main chain amino group of V47 and the carbonyl group of K52 of the D-loop hydrogen bond with the hydroxyl groups of Y145 and Y171, respectively (Figure 2B).

**Figure 2.**
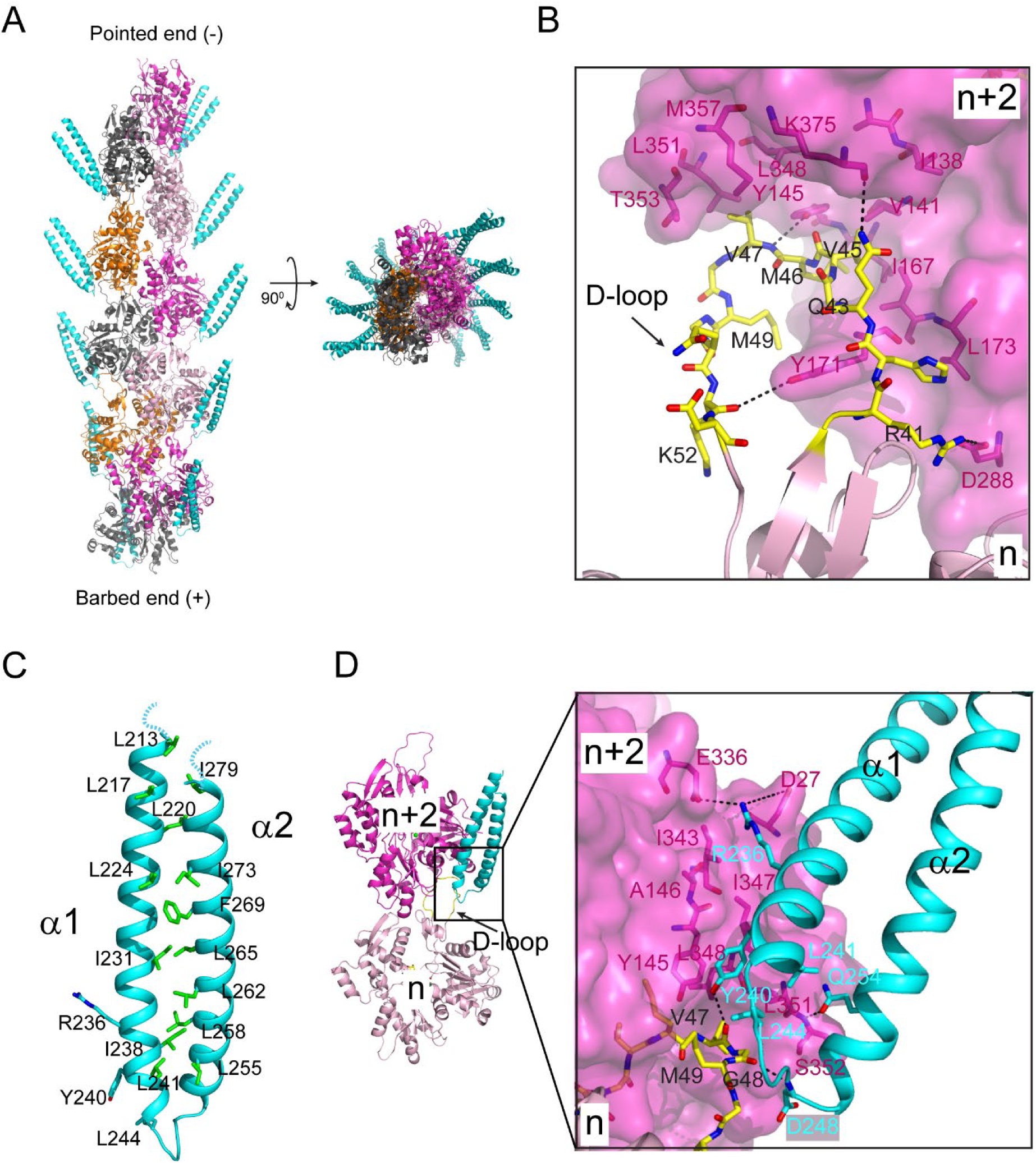
Cryo-EM structure of the Lfat1 ABD in complex with F-actin. (A) Cryo-EM structure of the F-actin-Lfat1 ABD complex. Left: side view of the structure positioned with the pointed end (-) up and barbed end (+) down. The visible part of ABD is colored in cyan. Right: Top view of the complex. (B) The D-loop conformation and its interactions in the hydrophobic cleft of the n+2^nd^ actin monomer. (C) Ribbon diagram of the distal portion of the Lfat1 coiled-coil domain. The two α-helices are zipped together through extensive hydrophobic interactions contributed mainly by leucine and iso-leucine residues (shown in sticks). (D) Structural representation of the interaction between the ABD domain of Lfat1 and F-actin. Inset: Extensive hydrophobic, hydrogen bonding, and electrostatic interactions were observed between the Lfat1 ABD domain and the two adjacent actin monomers (details see in the text).

The cryo-EM structure also revealed the distal portion of the coiled-coil hairpin, which consists of about 1/3 of the entire Lfat1 ABD domain (Figure S3D). The proximal end of the ABD domain is not visible, likely due to its flexibility, and zeroed out in 2D-class averaging. The structure revealed that the two α-helices of the Lfat1 ABD domain are zipped together by a stretch of hydrophobic residues, mostly leucines and isoleucines (Figure 2C). The ABD domain radiates away from the central F-actin core with its tip of the hairpin region binding to the site between two adjacent actin molecules within each strand of the F-actin filament (Figure 2A). The ABD domain embeds a surface area of 3703 Å^2^ on the actin filament, which is contributed by both the D-loop region of the n^th^ and the hydrophobic cleft of the n+2^nd^ actin monomers (Figure S4). Several hydrophobic residues (Y240, L241, and L244) located at the tip of the ABD hairpin are accommodated by a hydrophobic pocket formed between the two adjacent actin molecules (Figure 2D). The interaction between the ABD domain and the actin filament also involves several hydrogen bonds. The hydroxyl group of Y240 of the ABD domain forms a hydrogen bond with the main chain carbonyl group of V47 at the D-loop; the amino group of D248 of the ABD pairs with the carbonyl oxygen of G48; and the side chain carbonyl oxygen of ABD Q254 makes hydrogen bond with the main chain amine group of S352 of actin. Moreover, salt bridges are also observed between R236 of the ABD domain and D27 and E336 of the n+2^nd^ actin monomer (Figure 2D).

Together, our cryo-EM structure of the Lfat1 ABD domain in complex with F-actin revealed the intricate molecular basis of multivalent interactions between F-actin and a novel prokaryotic actin-binding domain. In addition, the complex structure revealed a 1:1 ratio of the interaction between the Lfat1 ABD and actin monomer, which is in agreement with the stoichiometry determined by the previous co-sedimentation experiment (Figure 2A, Figure S3C and D).

### Validation of key residues on the ABD domain in its recognition of F-actin

To validate our structural observations of the interaction between Lfat1 ABD and F-actin, we performed an alanine substitution mutagenesis experiment of three representative residues (R236, Y240, and Q254) in the Lfat1ABD domain (Figure 3A). We found that GFP-tagged R236A or Q254A ABD mutant showed a slight increase in diffused signals, with the majority of proteins remaining colocalized with F-actin. However, the Y240A mutation renders the protein mostly cytosolic (Figures 3B and C). To further validate the fluorescence imaging results, we performed an *in vitro* F-actin co-sedimentation titrating assay to measure the binding affinity between ABD proteins and F-actin (Figure 3D and E). The apparent *K_d_* for wild-type ABD to F-actin is calculated at about 1.48 µM, which is on par with LifeAct (*K_d_* of 2.2 µM) (Riedl *et al*, 2008). Consistently, the *K_d_* for R236A and Q254A mutants increased about 10-fold, 19.06 and 16.28 µM, respectively. More strikingly, the Y240A mutant showed a substantial decrease in affinity with a *K_d_* of 63.69 µM (Figure 3D and E). In summary, the mutagenesis experiments confirmed that multivalent interactions contribute to the binding of Lfat1 ABD with F-actin with hydrophobic interactions playing a central role, and the affinity and specificity were further enhanced by hydrogen bond and salt bridge interactions.

**Figure 3.**
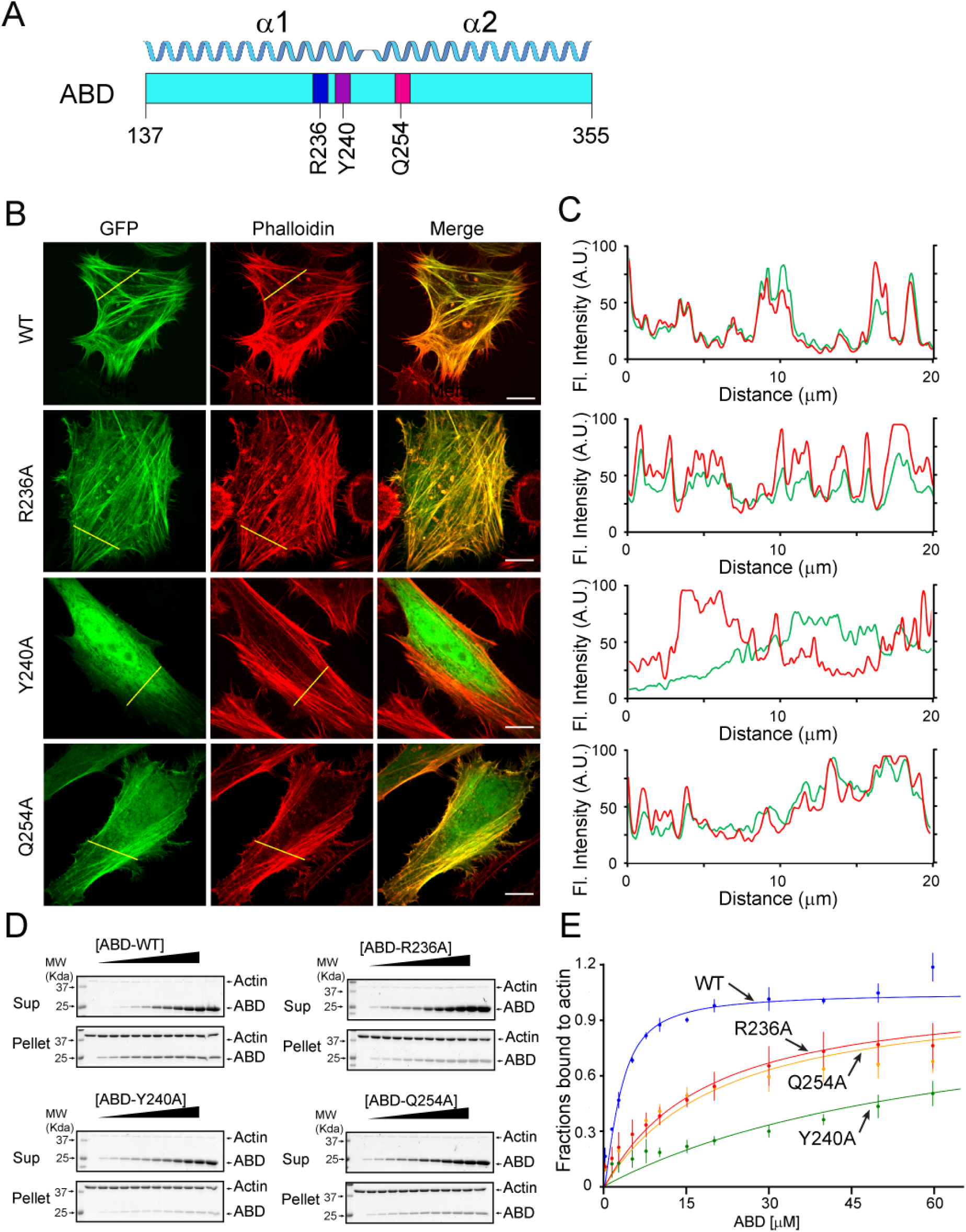
Validation of key Lfat1 ABD residues in their contributions to F-actin interactions. (A) Schematic diagram of the Lfat1 ABD domain. Three key residues (R236, Y240, and Q254) involved in actin binding are labeled. (B and C) F-actin localization analysis of indicated ABD mutants. GFP-Lfat1 WT, R236A, Y240A, or Q254A mutant was transiently expressed in HeLa cells followed by fixation and staining with phalloidin. Fluorescence images were taken by a confocal microscope and analyzed using Line scan along the indicated yellow lines. Scale bar = 10 µm. (D) Co-sedimentation assay of F-actin with WT and mutant Lfat1 ABD. Increasing amounts (0 to 60 µM) of recombinant WT or mutant ABD proteins were incubated with a fixed amount of actin. The samples were ultra-centrifugated after 30 min of room temperature incubation in 1x actin polymerization buffer. The supernatant and pellet fractions were analyzed by SDS-PAGE. (E) Quantitative analysis of the co-sedimentation titration data. The data point for each concentration was averaged from three independent experiments. The error bar represents the standard deviation.

### Comparison of Lfat1 ABD with other actin-binding domains

The discovery of a novel ABD from the *Legionella* effector Lfat1 prompted us to compare this unique ABD to other representative actin-binding proteins. Although ABDs adopt a diverse structural fold, most of them share a similar interaction scheme with F-actin by targeting a hotspot encompassing the D-Loop of Actin_n_ and the hydrophobic cleft of Actin_n+2_ (Figure 4A and B) (Dominguez, 2004). For example, LifeAct, which is derived from the first 17 residues from *Saccharomyces cerevisiae* ABP140 (Riedl *et al*., 2008), utilizes the hydrophobic residues aligned on one side of its amphipathic α-helix to engage primarily hydrophobic interactions with a small hydrophobic patch at the F-actin hotspot (Belyy *et al*, 2020; Kumari *et al*, 2020). The actin-binding CH1 domain of Utrophin (a neuromuscular junction scaffolding protein) contains multiple F-actin binding sites (Keep, 2000; Keep *et al*, 1999). Two of the actin-binding sites on the CH1 domain interact primarily with the D-loop region of the n^th^ actin monomer, and the third one, consisting of the N-terminal α-helix, spills the interface further into the subdomain I region of the n^th^ actin subunit (Kumari *et al*., 2020). The *Pseudomonas aeruginosa* effector protein, Exotoxin Y or ExoY, uses its C-terminal “anchor” helix to engage primarily hydrophobic interactions with the hydrophobic cleft in subdomain 1 of the n+2^nd^ actin subunit (Belyy *et al*, 2021) in a way similar to that of LifeAct (Belyy *et al*., 2020). ExoY also contains a peptide, which meanders on the surface of the n+2^nd^ actin subunit and functions as a “sensor” for the actin activator but contributes little affinity to F-actin binding (Belyy *et al*., 2021). These examples support that the hydrophobic cleft formed by subdomain 1 and 3 is the “hotspot” for many ABDs, and the binding site on actin is frequently extended to the vicinity of the “hotspot” depending on unique features associated with each ABD (Figure 4A).

**Figure 4.**
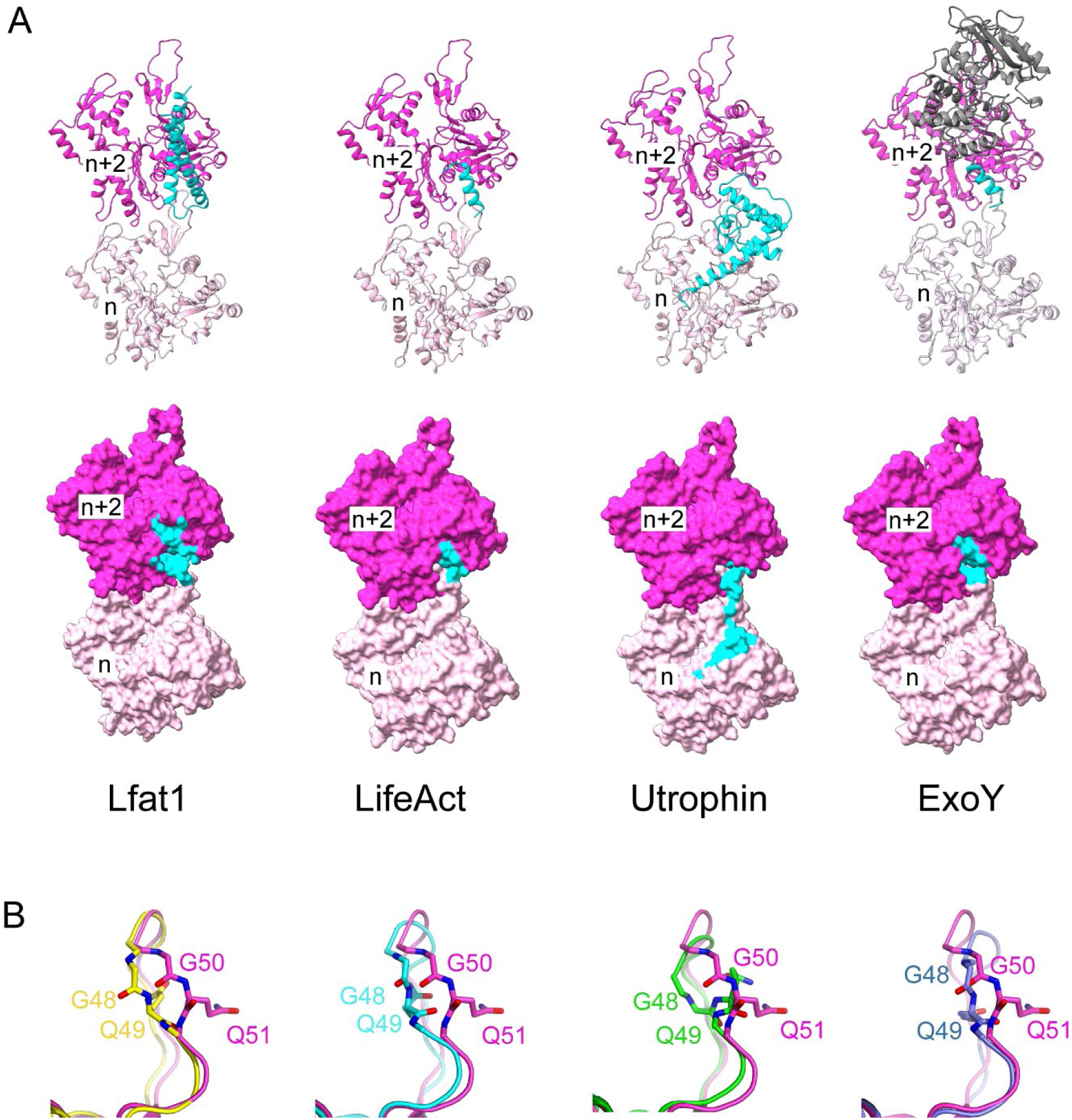
Structural comparison between Lfat1 and other ABD-F-actin complexes. (A) Ribbon diagram (Upper row) and surface representation (bottom row) of two adjacent actin monomers (purple and pink) bound with Lfat1 ABD, LifeAct (PDB ID: 7BTE), Utrophin (6M5G), and ExoY (7P1G). The region interfacing with each ABD is colored in cyan. (B) Structural comparison of the D-loop conformation in the Lfat1 ABD-F-actin complex (purple) with the D-loop in other F-actin-ABD complexes. F-actin alone (yellow), LifeAct (cyan), Utrophin (green), and ExoY (navy blue). Two D-loop residues with the largest deviation of backbone dihedral angles (G50 and Q51) are shown in sticks.

A structural comparison of the ABD-F-actin complex structures also revealed that the actin D-loop adopts a variety of conformations upon the binding of different ABDs (Figure 4B). In globular G-actin, the D-Loop is either disordered or adopts an α-helix, depending on its nucleotide state or binding with G-actin binding proteins (Graceffa & Dominguez, 2003). In F-actin, the D-Loop inserts itself into the hydrophobic target-binding cleft of the n+2 subunit immediately above it (Das *et al*, 2020; Dominguez & Holmes, 2011). The D-loop region is also involved in direct interactions with many ABDs (Figure 4). In the Lfat1 ABD and F-actin complex, specific hydrogen bonds are formed between the backbone carbonyl group of D-loop residues V47 and G48 and the ABD residues Y240 and D248 (Figure 2D). These hydrogen bonding interactions cause the D-loop to insert slightly deeper into the hydrophobic cleft and induce a unique conformation of the D-loop residues G50 and Q51 (*Bos taurus* numbering, equivalent to G48 and Q49 in *G. gallus*) not observed in other structures (Figure 4B). This observation suggests that although the hydrophobicity of the “hotspot” plays a dominant role in ABD-binding, the capacity to accommodate the large variety of actin-binding motifs at the “hot spot” is likely due to the structural plasticity of the D-loop.

### Engineering novel F-actin probes derived from the Lfat1 ABD

Fluorescent toxins or proteins are frequently used as F-actin probes in fixed or live cells, however, they all have certain limitations (Belin *et al*, 2014; Courtemanche *et al*, 2016; Lemieux *et al*, 2014; Munsie *et al*, 2009). The discovery of a new F-actin binding domain from the *Legionella* effector Lfat1 inspired us to investigate whether it can be developed as an alternative *in vivo* F-actin probe. We first tried to map the minimum F-actin binding region in Lfat1 ABD. A series of ABD truncations: ABD-S1 (residues 171-323), ABD-S2 (190-306), and ABD-S3 (211-280) were created and transiently expressed in Hela cells. All three truncated versions demonstrated specific colocalization with F-Actin comparable to the full-length ABD (Figure 5A). However, further shortening of this ABD resulted in loss of function and exhibited a complete cytosolic location (data not shown). Thus, our studies revealed a novel F-actin probe consisting of a 70 amino acid-long alpha-helix hairpin.

**Figure 5.**
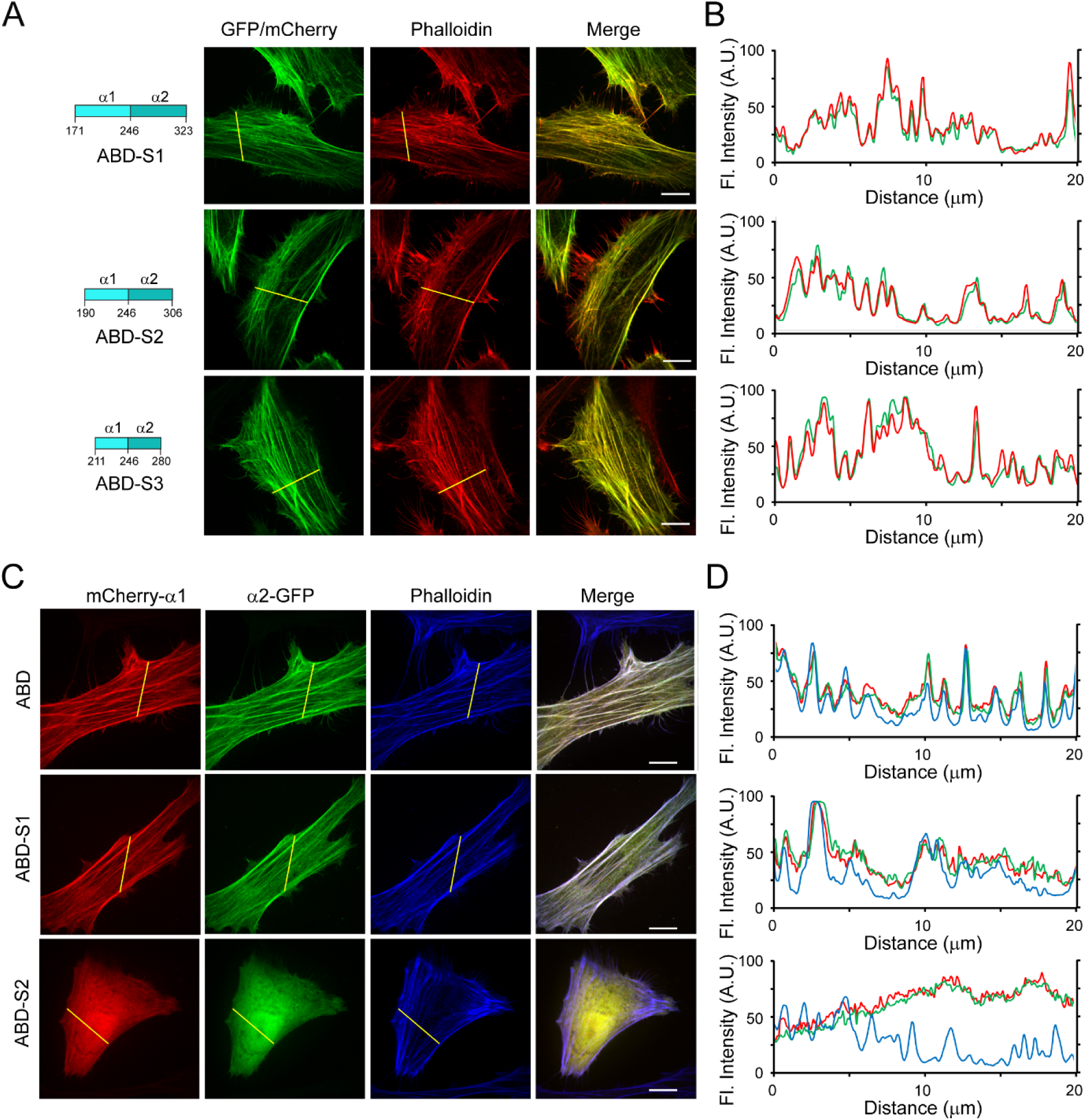
Engineering Lfat1 ABD as an *in vivo* F-actin probe. (A) Mapping the minimal actin-binding domain of Lfat1. Schematic of shortened Lfat1ABD fragments used for F-actin binding (Left). Representative fluorescence images of cells expressing indicated ABD fragments and stained with rhodamine-phalloidin. (B) Line-scan analysis for the indicated ABD probes along the yellow lines. (C) Representative fluorescence images of cells transiently transfected with plasmids expressing separated alpha helixes (mCherry-α1 and α2-GFP) of ABD, ABD-S1, and ABD-S2. The cells were fixed and stained with CF647-phalloidin. (D) Line-scan analysis of the images shown in (C). Scale bar = 10 µm.

The Lfat1 ABD contains two long alpha helixes forming a hairpin. We next asked whether this ABD remains functional if the alpha-helix hairpin is split into two individual alpha helixes. To test this, we fused an N-terminal mCherry with α1 and a C-terminal GFP with the α2 of the ABD, respectively, and examined their intracellular localization. All these single alpha helix fusions showed a diffused localization (Figure S5A, B), however, to our surprise, when the two fusion constructs encoding full-length ABD-α1 and α2 were expressed together, these two alpha helixes were able to form a functional ABD and colocalized with F-Actin as the intact wild type ABD (Figure 5C and D). Interestingly, the alpha helixes derived from ABD-S1 could also reconstitute a functional F-actin probe, but not the further shortened alpha helices derived from ABD-S2 (Figures 5C and D). These results suggested that the F-actin probe derived from the Lfat1 ABD can be used in a split form, providing flexibility to this new probe.

### Lfat1 is a lysine fatty acyltransferase (KFAT)

The AlphaFold-predicted structure of Lfat1 revealed a globular domain composed of the N-and C-terminus beside the central, coiled-coil hairpin (Figure 1A). Structural homology search using the DALI server (Holm & Rosenstrom, 2010) yielded the top hit as the RID (Rho GTPase Inactivation Domain) toxin from the *Vibrio vulnificus* (PDB:5XN7) with a Z-score of 7.9. The catalytic domain of both proteins contains a central β-sheet flanked by multiple α helices. The conserved catalytic dyad (H38 and C403) in Lfat1 were positioned with a similar orientation to the dyad in the RID toxin (H2595 and C2835) (Figure 6A).

**Figure 6.**
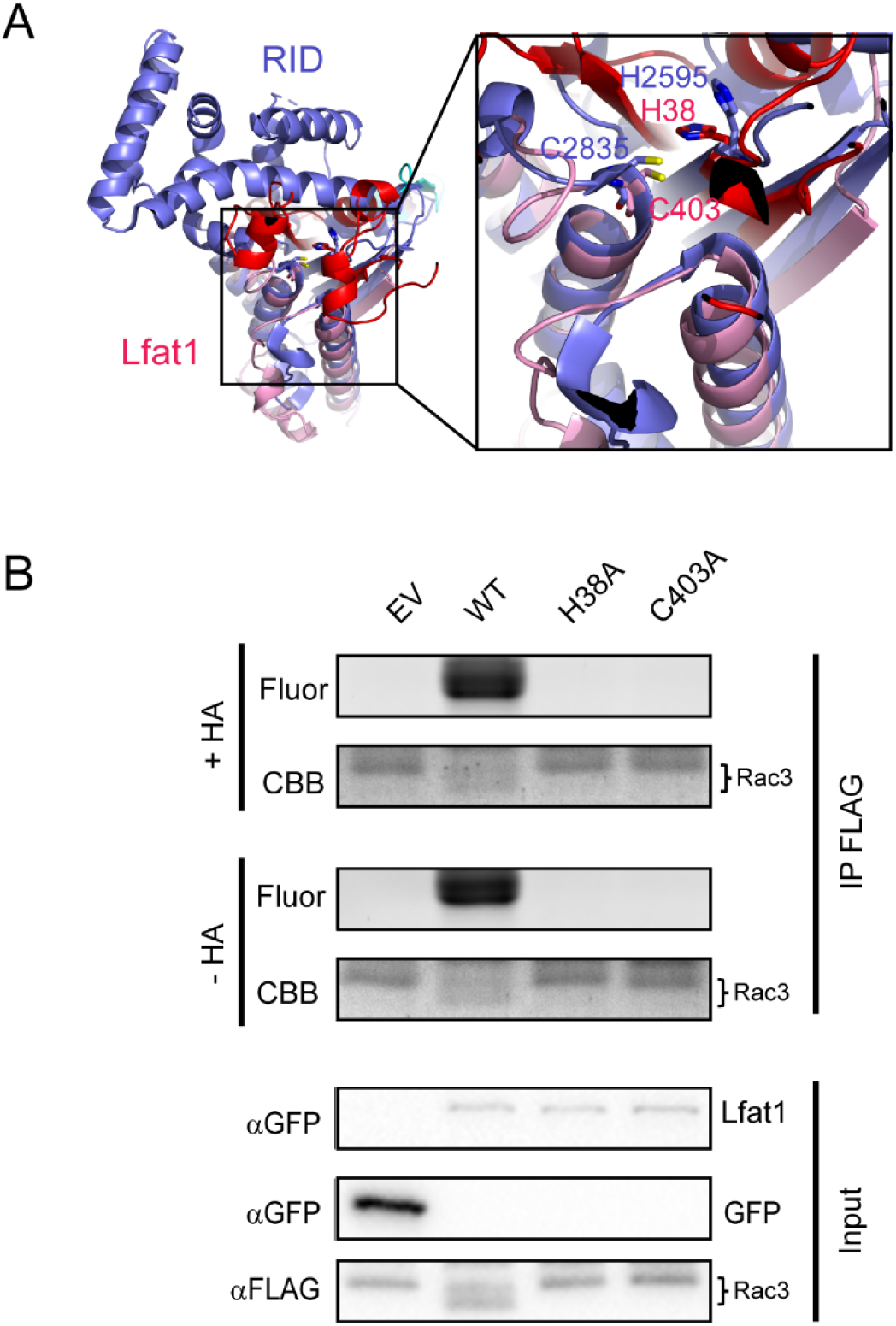
Lfat1 is a lysine fatty acyltransferase that modifies eukaryotic small GTPases. (A) Ribbon representation of the *Vibrio vulnificus* RID (PDB ID: 5XN7) catalytic domain (purple) superimposed with the AlphaFold-predicted NC-domain of Lfat1 (N-terminal domain in red, C-terminal domain in pink). Inset: the catalytic pockets of the two proteins with conservation of the catalytic histidine and cysteine between RID and Lfat1 shown in sticks. (B) Lfat1 catalyzes lysine fatty acylation of Rac3. N-terminal Flag-tagged Rac3 was co-expressed with either GFP empty vector (EV), GFP-Lfat1 WT, H38A, or C403A mutant in HEK293T cells for 24 hours. Flag-Rac3 was enriched using immunoprecipitation and subjected to Click chemistry. The samples were then separated on SDS-PAGE and scanned for fluorescence signals.

The RID toxin is a lysine fatty acyltransferase (KFAT) processed from a much larger prototoxin that transfers long acyl chains to the ε-amine of lysines in small GTPases, such as those in the Rac subfamily (Zhou *et al*, 2017). To determine whether Lfat1 possesses a similar KFAT activity, we probed the fatty acylation of a small GTPase Rac3. Briefly, HEK-293T cells were co-transfected with plasmids expressing Flag-Rac3 and GFP-Lfat1 or its catalytic mutants followed by treatment with Alk14, a clickable chemical analog of palmitic acid modified with a terminal alkyne moiety. Flag-Rac3 proteins were enriched from transfected cells by immunoprecipitation and were then conjugated to an azide-containing fluorophore, TAMRA-N_3_ via copper-catalyzed cycloaddition (Hein *et al*, 2008). The reaction products were separated by SDS-PAGE and analyzed by in-gel fluorescence detection and Western blot. Robust fluorescence signals associated with Rac3 were detected in the presence of wild-type Lfat1, but not its H38A or C403A catalytic mutants (Figure 6B). Furthermore, we found that Rac3-associated fluorescence signals were stable even after the hydroxylamine (HA) treatment, which can reverse acylation on cysteine but not lysine residues. We next asked whether Lfat1 can fattyacylate other host targets. To address this question, we performed a click chemistry-coupled SILAC (Stable Isotope Labeling by Amino Acids in Cell Culture) mass-spectrometry experiment to identify potential targets (Figure S6A). Interestingly, nearly half of the top hits are small GTPases, including Rabs, RheB, RalA, and Rap1B (Figure S6B). Fattyacylation by Lfat1 on several of the small GTPases was further verified using click chemistry (Figure S6C-F). Furthermore, the fatty acylation of small GTPases by Lfat1 does not appear to depend on F-actin binding, as these substrates were still modified by the actin-binding-deficient Lfat1 Y240A mutant (Figures S7A and B). Together, our results demonstrated that the *Legionella* effector Lfat1 is a *bona fide* KFAT that potentially fattyacylates host small GTPases when exogenously expressed in cultured cells.

## Discussion

In this study, we identified a novel actin-binding domain from *Leginella pneumophila*. This prokaryote-originated ABD has an alpha-helical hairpin-like structure, which is unprecedented from any other known ABDs. We further mapped the minimum region (∼ 70 residues) required for actin binding and demonstrated the feasibility of using this prokaryote ABD as an alternative F-actin probe. Another unique and potentially useful characteristic of this probe is its ability to function as a split-ABD. Although the individual alpha helix derived from the Lfat1 ABD fails to bind actin, they were able to reconstitute a functional intact ABD when co-expressed. This unique feature can be harnessed to target multi-component biological complexes to F-actin by genetically fusing individual components to the split alpha helixes. Furthermore, our cryo-EM structure revealed that the Lfat1 ABD intersects with F-actin obliquely. Its N- and C-termini are pointed away with an adjustable distance from the filament depending on the size of the designed ABD. Thus, the Lfat1 ABD can be used to target proteins of interest to F-actin with a tunable distance from the filament to achieve spatial distribution-related specificity.

F-actin probes are essential tools in cell biology for visualizing and studying the dynamics of the actin cytoskeleton in living and fixed cells. These probes come in various forms, including fluorescently labeled phalloidins, actin-binding proteins, and genetically encoded fluorescent actin markers (Melak *et al*, 2017). Each type of probe has its advantages and limitations. The choice of probe depends on the specific experimental requirements, such as whether live-cell imaging is needed, the level of perturbation that can be tolerated, and the ease of use. The toxic chemical derived from fungi, phalloidin, has been developed as the gold standard F-actin marker to stain actin in fixed samples and tissues (Cooper, 1987). However, it is not suitable for live-cell imaging due to its toxicity and low cell permeability. Many yeast- or human-derived actin-binding domains have been developed into F-actin probes by fusion with fluorescent proteins. Lifeact (Riedl *et al*., 2008), utrophin (Burkel *et al*, 2007), and F-tractin (Brehm *et al*, 2004) are the three most commonly used genetically encoded probes. Although these probes have been widely used for live-cell imaging, they suffer from problems such as low affinity for F-actin and perturbation in actin dynamics. Our discovery of a novel ABD offers an alternative F-actin probe that not only can be used to study actin dynamics but also can be used to as a versatile anchor to target specific activities to F-actin.

The discovery of Lfat1 as an F-actin–binding lysine fatty acyl transferase raised the intriguing question of whether its enzymatic activity depends on F-actin binding. Recent studies have shown that other *Legionella* effectors, such as LnaB and Ceg14, use actin as a co-factor to regulate their activities. For instance, LnaB binds monomeric G-actin to enhance its phosphoryl-AMPylase activity toward phosphorylated residues, resulting in unique ADPylation modifications in host proteins (Fu *et al*, 2024; Wang *et al*, 2024). Similarly, Ceg14 is activated by host actin to convert ATP and dATP into adenosine and deoxyadenosine monophosphate, thereby modulating ATP levels in *L. pneumophila*–infected cells (He *et al*, 2025). However, this does not appear to be the case for Lfat1. We found that Lfat1 mutants defective in F-actin binding retained the ability to modify host small GTPases when expressed in cells (Figure S7). These findings suggest that, rather than serving as a co-factor, F-actin may serve to localize Lfat1 via its actin-binding domain (ABD), thereby confining its activity to regions enriched in F-actin and enabling spatial specificity in the modification of host targets.

Our finding that Lfat1 is a protein lysine fatty acyl transferase provides important insights to understand the physiolical function of Lfat1. The Rho Inactivation Domain (RID) is a module found in Multifunctional-Autoprocessing Repeats-in-Toxin (MARTX) toxins produced by certain Gram-negative pathogenic bacteria, such as *Vibrio cholerae* and *Vibrio vulnificus* (Satchell, 2015). RID primarily targets the Rho GTPase family members by the covalent attachment of long-chain fatty acids to the ε-amino groups of lysine residues. The modification inactivates the small GTPases, leading to disruption of the actin cytoskeleton and consequent cell rounding and hence facilitating bacterial invasion and impairing host immune responses (Zhou *et al*., 2017). A recent study reported that RID modifies other host proteins, notably septins. Fatty-acylation on septins alters the localization and compromises the host cell structural integrity (Xu *et al*, 2024). In this study, we demonstrated that the globular NC-domain of Lfat1 exhibits KFAT activity when overexpressed *in vivo*. Using Click chemistry, we showed that Lfat1 could fatty-acylate lysines of the host small GTPase Rac3 (Figure 6B) and other small GTPases (Figure S6). Many of the small GTPase substrates we identified, are known to associate with and regulating actin. For example, RhoG regulates the actin cytoskeleton in lymphocytes (Vigorito *et al*, 2003); Rap1 is reported to regulate actin reorganization and microtuble organizing center polarization at the B cell immune synapse (Wang *et al*, 2017); RheB is reported to regulate actin filament distribution (Gau *et al*, 2005); Ral GTPases (RalA and RalB) links Ras, Rac, Rho signaling to control cell migration (Zago *et al*, 2019); and RAB8A regulates spindle migration via ROCK-mediated actin assembly in mouse oocyte meiosis (Pan *et al*, 2019). The identification of the KFAT effectors among all *Legionella* species and set up a solid foundation for further characterizations of this family of effectors. However, future studies will be needed to identify which substrates are physiologicaly important under infection conditions.

## Acknowledgments

We thank Dr. Sheng Zhang and Dr. Qin Fu for helping with the proteomics study. W.Z. acknowledges support from the T32 training grant and the Sadov Graduate Student Fellowship (Cornell University). This work is supported by the National Institutes of Health R01GM144452(Y.M.), R01AI153110 (H.L.), and HHMI.

## Material and methods

### Plasmid construction

Please refer to Supplemental Table 2 for constructs used in this study. All targeted DNA fragments were amplified by PCR using the primers listed in Supplemental Table 3. The amplified DNA fragments were digested by restriction enzymes, BamHI and XhoI, and ligated to corresponding vector plasmids. Plasmids containing the targeted genes were amplified using NEB^®^ Stable Competent C3040I *E. coli* strain and verified by sequencing. For mutagenesis, plasmids containing wild-type genes were used as templates with mutagenic primers in a PrimeStar MasterMix PCR reaction to amplify the entire plasmid, followed by DpnI digestion, and plasmids carrying the desired mutations were amplified using the C3040I *E. coli* strain and verified by sequencing.

### Protein purification

Actin was purified from muscle acetone powder prepared from ground beef (Pardee & Spudich, 1982). Briefly, 10 grams of muscle acetone powder was dissolved with 200 ml G-actin buffer (5 mM Tris pH 7.5, 0.2 mM CaCl_2_, 0.2 mM ATP, 0.5 mM DTT) at 4 °C with stirring for 30 minutes followed by low-speed centrifugation at 5000-10,000g for 10-20 minutes to remove insoluble solids. Soluble extracts were centrifuged at 15,000g for 60 min at 4 °C, and actin was precipitated by the slow addition of solid ammonium sulfate to reach 25% saturation. Precipitates were collected by centrifugation at 15,000g for 30 min and dissolved in 50 ml G-actin buffer and dialyzed against 2 liters of G-actin buffer overnight with two changes of buffer. The dialyzed actin solution was centrifuged at 32,000g for 60 min to remove any aggregates. Actin was polymerized by the addition of 10x polymerization buffer to a final solution containing 150 mM KCl, 2 mM MgCl_2_, 2 mM EGTA, and 1 mM ATP. Polymerization was allowed to proceed for 30 min at 25 °C, then in a cold room for 90 min. F-actin was pelleted by centrifugating at 100,000g for 30 min, and the pellet was resuspended in 10–20 ml G-actin buffer and homogenized in a glass-glass homogenizer on ice. The suspension was dialyzed against 2 liters of G-actin buffer for 48 hours with four changes of dialysis solution to completely depolymerize the F-actin. The solution was further clarified by centrifugation at 32,000g for 60 min. The clarified supernatant was loaded onto a Sephadex G-150 or Sephacryl S-100 HR column, equilibrated in G-actin buffer, to remove the remaining contaminating G-actin-binding proteins. Peak fractions were analyzed by SDS-PAGE, pooled, and flash-frozen in liquid nitrogen before storage.

Recombinant Lfat1 ABD (WT and mutants) proteins were expressed in Rosetta *E. coli*. The expression was induced by 0.1 mM IPTG at 18 °C overnight. The bacteria were harvested by centrifugation at 4,000 RPM using the Beckman Coulter JLA-9.1000 rotor for 20 minutes. The cell pellets were lysed in Buffer A (20 mM Tris pH7.5, 150 mM NaCl) using sonication. The whole cell lysate was then centrifuged at 16,000 RPM using the Beckman Coulter JA-25.50 rotor for 40 minutes at 4 °C, and the clarified lysate was then bound to Cobalt resins on a rotator at 4 °C for 2 hours. The resins were then extensively washed with Buffer A, and the ABD proteins were released from the resin by a SUMO-specific protease, Ulp1. The released proteins were collected and further purified using size-exclusion chromatography on a Superdex S75 column. The purified ABD proteins were concentrated and then flash-frozen in liquid nitrogen for storage.

### Cell culture and co-immunoprecipitation

HEK293T and HeLa cells were maintained at a low passage and grown in DMEM (Dulbecco’s Modified Eagle Medium) supplemented with 10% FBS (fetal bovine sera). For transfection, plasmids were mixed with 1 mg/ml PEI (polyethylenimine MW10K, Millipore Sigma) at a 5:1 ratio in DMEM incubated for 15 minutes at room temperature, and added directly to cells. After 24 hours of transfection, the cells were used either for immunoprecipitation or imaging experiments.

For immunoprecipitation, HEK293T cells transfected with the indicated plasmids after 24 hours were chilled on ice, washed with ice-cold PBS, and detached using lysis buffer (20 mM Tris pH7.5, 150 mM NaCl, 1 mM DTT, 0.5% Triton X-100, 0.1% sodium deoxycholate, 1 mM PMSF, with Roche protease inhibitor cocktail). The cells were then lysed using a sonicator on ice. The lysate was clarified by centrifugation at 16,000 RPM using the Beckman Coulter JA-25.50 rotor for 15 minutes at 4 °C. The lysate was then incubated with anti-GFP nanobody beads for 2 hours with rotating. The mixture was then washed in a buffer containing 20 mM Tris pH7.5, 150 mM NaCl, 1 mM DTT, and 0.5% Triton X-100. The washed beads were then dissolved in 1X SDS loading buffer, and the samples were analyzed by SDS-PAGE followed by Western-blot and scanned using Li-COR Odyssey CLx scanner.

### Fluorescence Microscopy

HeLa cells grown on glass cover slides were transfected with the indicated plasmids after 24 hours. The cells were then washed with PBS and fixed in 4% PFA in PBS for 15 minutes at room temperature. The fixed cells were washed twice with PBS and incubated with Odyssey blocking buffer with 0.1% saponin and rhodamine- or CF647-phalloidin (Thermofisher) for 1 hour at room temperature. The stained cells were washed three times with PBS and mounted on a glass specimen slide with Fluoromount-G (Thermofisher), and were imaged using a 3i spinning-disc confocal fluorescence microscope. Line-scan analysis was performed using ImageJ (Schneider *et al*, 2012).

### CryoEM sample preparation, data collection, data processing

Purified G-actin (1 mg/ml) was mixed with an equal molar of ABD in G-actin buffer and incubated overnight at 4 °C. The ABD-F-actin complex samples with a serial dilution were applied to a glow-discharged copper Quantifoil r1.2/1.3 grids and rapidly plunged frozen in liquid ethane using FEI-Vitrobot-Mark-IV. The vitrified grids were then transferred to liquid nitrogen for storage and data collection.

For data collection, the grids were imaged using a ThermoFisher Talos Arctica 200 kV electron microscope with a K3 direct electron detector and a Gatan bioquantum energy filter and the dataset was collected using SerialEM software using the following parameters: -0.4 to -3.0 µm defocus range, pixel size of 1.5879062 Å, total electron dose of 40.68 electrons per Å^2^, exposure time of 1.23 seconds, 50 frames per movie, and 4548 total movies. The movies were then imported into CryoSPARC (Punjani *et al*., 2017) for motion correction and patch-CTF estimation. 2173 out of 4548 micrographs were selected after manual curation. The CryoSPARC Helical Tracer job was utilized to perform particle picking with a minimum particle diameter of 50 Å and a maximum diameter of 100 Å, with each particle separated by 100 Å. A total of 1,733,108 particles were extracted from micrographs using a box size of 480 pixels. Iterative 2D classifications were performed on these particles, and low-resolution classes were discarded after each iteration.

Finally, 1,220,462 particles were used for ab-initio initial model building, followed by iterative helical and local refinements. Final map was sharpened using CryoSPARC’s Sharpening Tool using half maps and a B-factor of 58.87 as obtained from the Guinier plot.

### Model building, refinement, validation

The atomic model of F-actin (PDB: 7BTI) and the AlphaFold-predicted model of Lfat1ABD were docked into the finalized CryoEM density using ChimeraX 1.25 (Pettersen *et al*, 2021). The atomic model of the F-actin-ABD complex was then refined iteratively using Phenix (Adams *et al*, 2010), and the final model, CryoEM full-map and half-maps, were validated by the Worldwide Protein Data Bank (wwPDB).

### F-actin co-sedimentation assay

Recombinant Lfat1 ABD was mixed with buffer control or G-actin at a final concentration of 20 µM. The mixed samples were either incubated in G-actin buffer or F-actin polymerization buffer (50 mM KCl, 2 mM MgCl2) at room temperature for 30 minutes followed by centrifugation at 70,000 RPM using the Beckman Coulter TLA-100.3 rotor for 30 minutes at 4 °C. The supernatant and pellet were then analyzed on SDS-PAGE.

To determine the F-actin binding affinity of ABD, purified recombinant Lfat1 ABD WT and mutant proteins were diluted in G-actin buffer in a series of concentration at 0, 1.25, 2.5, 5, 7.5, 10, 15, 20, 30, 40, 50, 60 µM and were incubated with 20 µM of G-actin. Actin polymerization was initiated at room temperature for 30 minutes by adding F-actin polymerization buffer. The samples were then centrifugated at 70,000 RPM using the Beckman Coulter TLA-100.3 rotor for 30 minutes at 4 °C, and the supernatants and pellets were then analyzed on SDS-PAGE and staining Coomas Brilliant Blue dye, and the intensity of the bands was quantified using ImageJ. and the intensity of the ABD band was divided by the intensity of the actin band in the pellet fractions to calculate the % of ABD bound to F-actin. The average of each data point from three independent experiments was then plotted, and an exponential fit was used to calculate the apparent K_d_ in R studio.

### Protein fatty-acylation detection via Click chemistry

Indicated plasmids were transfected into HEK 293T cells using PEI transfection reagent. After overnight transfection, cells were treated with 50 μM Alk14 (Cayman Chemical) for 6 hours. The cells were washed with ice-cold PBS and then lysed in lysis buffer (25 mM Tris-HCl, pH 7.8, 150 mM NaCl, 10% glycerol, and 1% NP-40) with protease inhibitor cocktail at 4 °C for 30 min. After centrifugation at 17,000 g for 30 min at 4 °C, the supernatant was collected and incubated with 20 μL of anti-Flag affinity beads (Sigma Aldrich) at 4 °C for 2 h. The affinity beads was washed three times with washing buffer (25 mM Tris-HCl, pH 7.8, 150 mM NaCl, 0.2% NP-40) and resuspended in 20 μL of washing buffer. TAMRA-N3 (Lumiprobe), TBTA (TCI chemicals), CuSO_4_, and TCEP (Millipore) were added into the reaction mixture in the order listed. The click chemistry reaction was allowed to proceed at room temperature for 30 min. The reaction was quenched by adding 6x SDS loading dye and boiled for 5 min. Where indicated, samples were treated with hydroxylamine (HA) to remove cysteine fatty acylation. The samples were then separated on SDS-PAGE and fixed in a buffer (50% CH_3_OH, 40% water and 10% acetic acid) by shaking for 1 hours at 4°C and then washed and store in water. The gel was scanned to record fluorescence signal using a ChemiDoc MP (BioRad).

### SILAC sample preparation and MS data analysis

SILAC samples were prepared from cells transiently expressing WT or H38A mutant Lfat1 using a published protocol (Kosciuk *et al*, 2020). The samples were then trypsin digested and the peptides were analyzed using an Orbitrap Fusion^TM^ Tribrid^TM^ (Thermo-Fisher Scientific) mass spectrometer. The MS and MS/MS spectra were subjected to database searches using Proteome Discoverer (PD) 2.4 software (Thermo Fisher Scientific, Bremen, Germany) with the Sequest HT algorithm. The database search was conducted against a *Homo sapiens* Uniprot database with the following variable modifications: methionine oxidation; deamidation of asparagine/glutamine; SILAC heavy: R10 (10.008 Da) and K8 (8.014 Da) and light labeling on R and K; palmitoylation plus biotin on K, protein N-terminus and fixed modification of cysteine carbamidomethylation.

---

**Table 1.** CryoEM Data collection, refinement and validation statistics.

|  |  |
| --- | --- |
| **Microscope** | FEI Talos Artica |
| **Voltage (keV)** | 200 |
| **Defocus range (um)** | -0.4 to -3.0 |
| **Camera** | K3 direct electron detector |
| **Pixel size (Å)** | 0.833 (super-resolution) |
| **Total electron dose (e/Å<sup>2</sup>)** | 40.68 |
| **Exposure time (seconds)** | 1.23 |
| **Frames per movie** | 50 |
| **Number of images** | 4548 |
| **3-D refinement statistics and helical symmetry** |  |
| *Total number of particles* | 1,220,462 |
| *Resolution (Å)* | 3.58 |
| *Helical twist* | -167 |
| *Rise* | 28 |
| **Model composition and validation** |  |
| *Non-hydrogen atoms* | 34,740 |
| *Protein residues* | 4390 |
| *Ligands* | 10Mg, 10ADP |
| **RMSD** |  |
| *Bond lengths(Å)* | 0.25 |
| *Bond angles (°)* | 0.5 |
| *B-factor (Å<sup>2</sup>) Protein* | 58.87 |
| *B-factor (Å<sup>2</sup>) Ligand (ADP)* | 56.6 |
| **MolProbity Score** | 1.9 |
| **Clashscore** | 5 |
| **Ramachandran plot:-Favored** | 95 |
| **Allowed** | 5 |
| **Outlier** | 0 |
| **PDB ID** | 8VAA |
| **EMDB Code** | 43087 |

**Figure 1 – Figure supplement 1.**
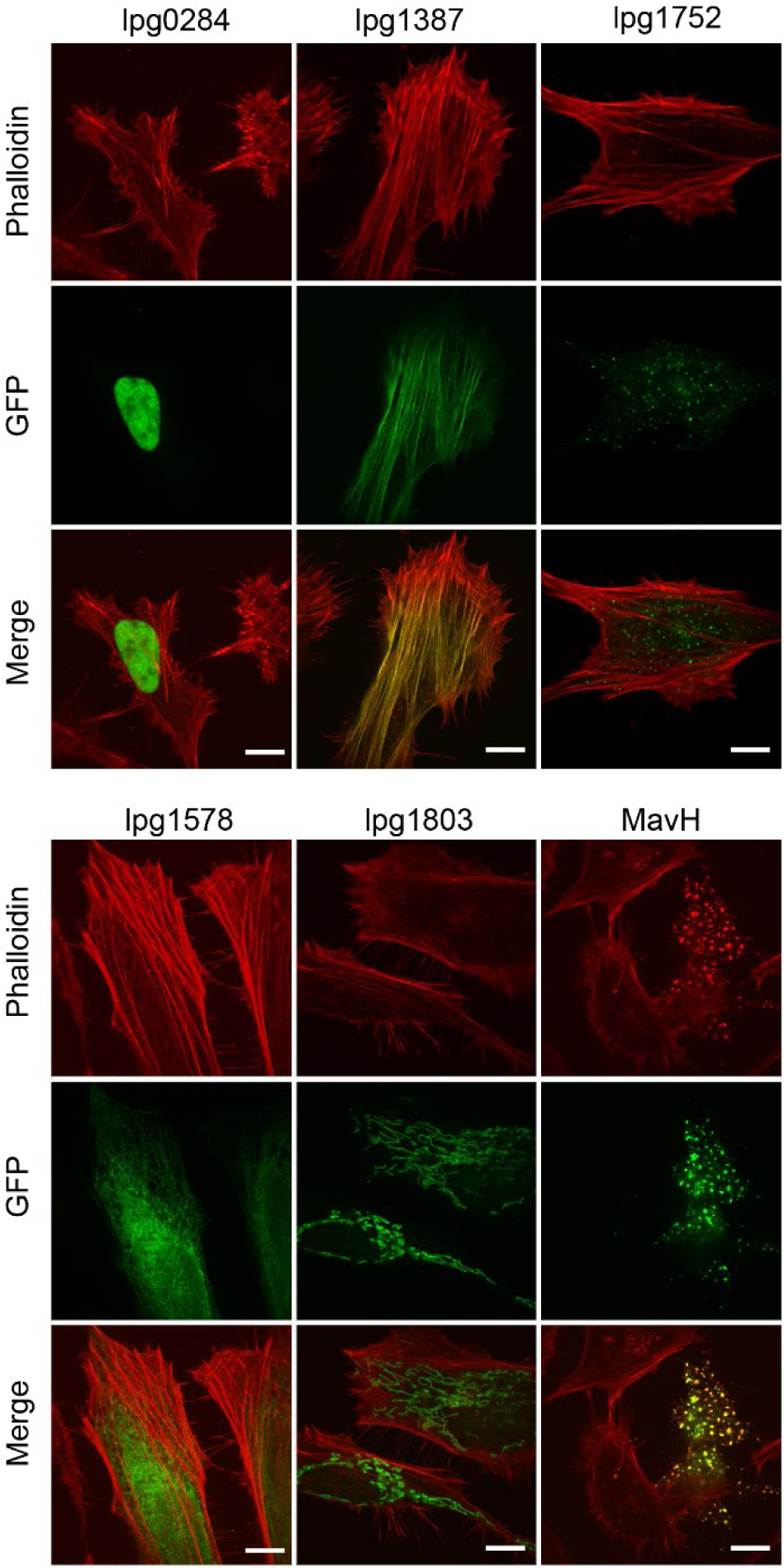
Intracellular localization representative Legionella effectors. Individual plasmid from a library consisting of 315 GFP-tagged *Legionella pneumophila* effectors was transiently expressed in HeLa cells followed by PFA fixation, stained with rhodamine-phalloidin, and visualized by fluorescence microscopy. Images showing intracellular localization of several representative effectors: lpg0284 (nuclear); lpg1387 and MavH (F-actin); lpg1578 (ER); and lpg1803 (mitochondrial). Scale bar = 10 µm.

**Figure 2 – Figure supplement 1.**
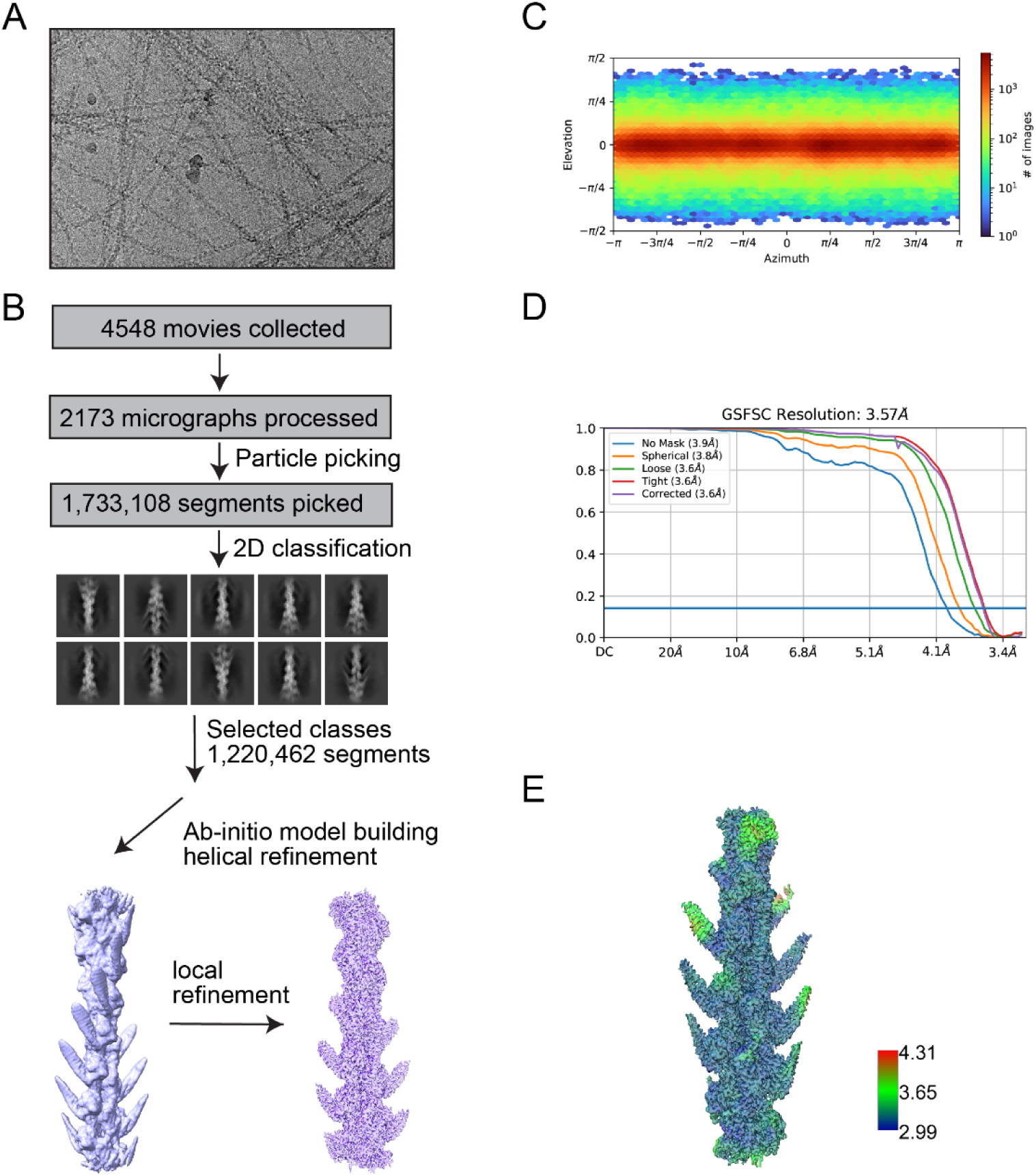
CryoEM data processing details of F-actin-Lfat1ABD complex. (A) Representative CryoEM micrograph of Lfat1ABD-F-actin complex. (B) Data-processing and structure determination workflow. Movies were collected at Talos Arctica electron microscope. Motion correction and patch CTF estimation were performed using CryoSPARC. Following manual curation, automatic filament tracer-based particle picking, and extraction, 1.2 million particles were 2-D classified and an ab initio model was built and refined using helical refinement until convergence. (C) Angular distribution of particles. (D) Resolution estimation by GSFSC. (E) Local resolution map of the final CryoEM map.

**Figure 2 – Figure supplement 2.**
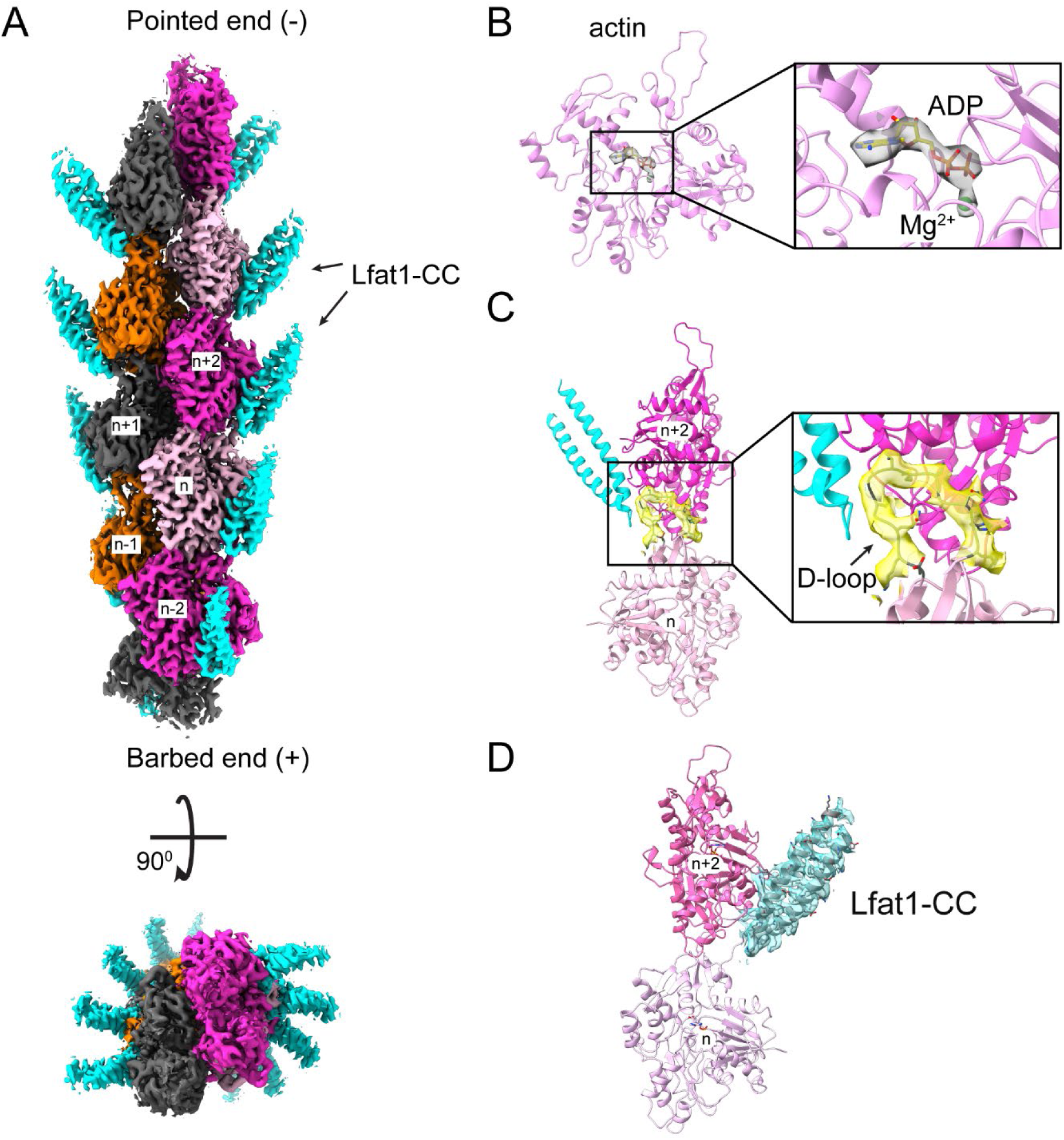
CryoEM maps of the F-actin-Lfat1ABD complex. (A) Final CryoEM density map of the complex. Top: Side view of the complex density map with the pointed end (-) up and barbed end (+) down. Lfat1 ABD is shown in cyan. Bottom: Top view of the complex density map. (B) The EM density map (grey) of ADP-Mg^2+^ at the ATP binding cleft of actin subunits. (C) The EM-map of actin D-loop (yellow). (D) The EM density map (cyan) of the distal region of the Lfat1 hairpin.

**Figure 2 – Figure supplement 3.**
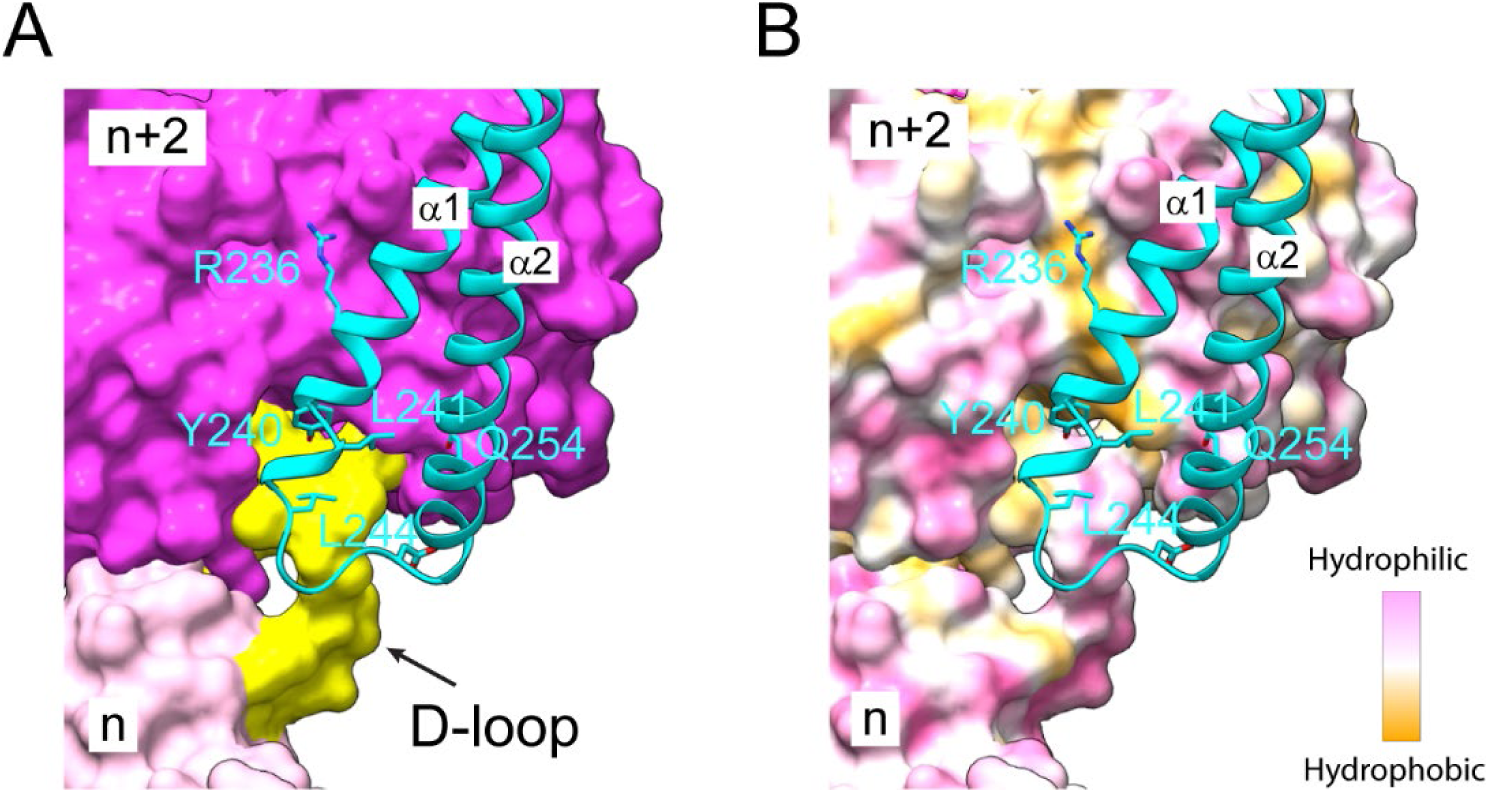
The interface between Lfat1 ABD and F-actin. (A) Surface representative of two adjacent actin molecules (purple and pink) and the D-loop region of the n^th^ actin molecule is colored yellow. Lfat1 ABD is represented as a ribbon in cyan. Key residues involved in F-actin interaction are shown in sticks. (B) The interface between Lfat1 ABD and F-actin at the same orientation as in (A). The surface of the two actin molecules is colored based on hydrophobicity, with hydrophobic areas in orange and hydrophobic regions in pink.

**Figure 5 – Figure supplement 1.**
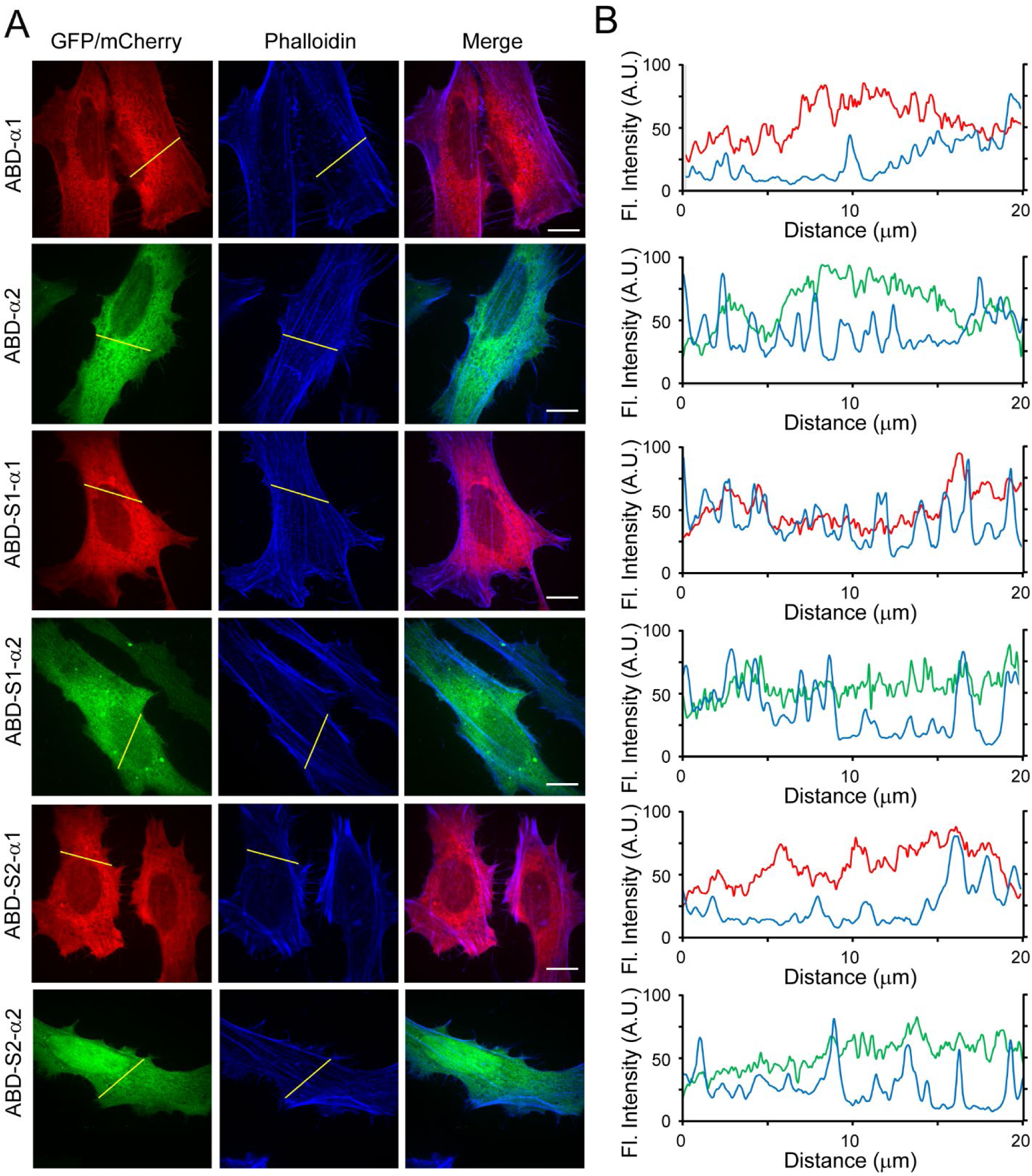
Intracellular localization of separated individual alpha helixes of the Lfat1 ABD. (A) Representative images of HeLa cells transiently expressing individual alpha-helixes from full-length and truncated Lfat1 ABD. Cells were fixed and stained with CF647-phalloidin. (B) Line-scan analysis of constructs tested in (A). Scale bar = 10 µm.

**Figure 6 – Figure supplement 1.**
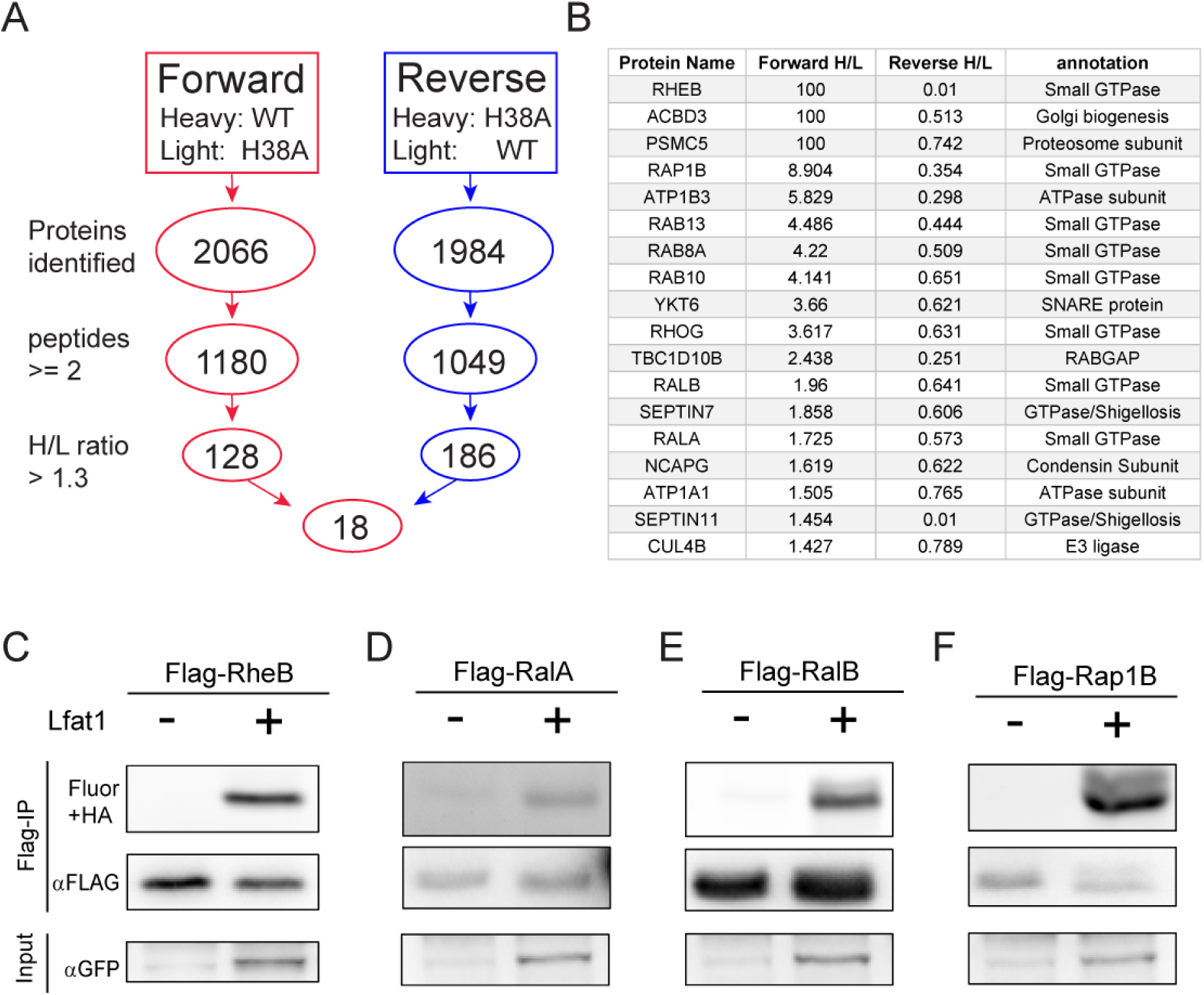
Identification of potential Lfat1 substrates by Click chemistry-coupled SILAC mass spectrometry. (A) Flow chart for selecting high-confidence substrate hits from the SILAC screen. (B) Top hits of potential Lfat1 substrates identified in this SILAC-MS experiment. (C) – (F) Click chemistry verification of some small GTPases selected from the top hit list. N-terminal Flag-tagged RheB, RalA, RalB, or Rap1B was co-expressed along with GFP-Lfat1 or GFP control in HEK293T cells followed by anti-Flag immunoprecipitation and Click chemistry conjugation reaction with an Azide-containing fluorophore. The samples were separated on SDS-PAGE and scanned for fluorescence signals.

**Figure 6 – Figure supplement 2.**
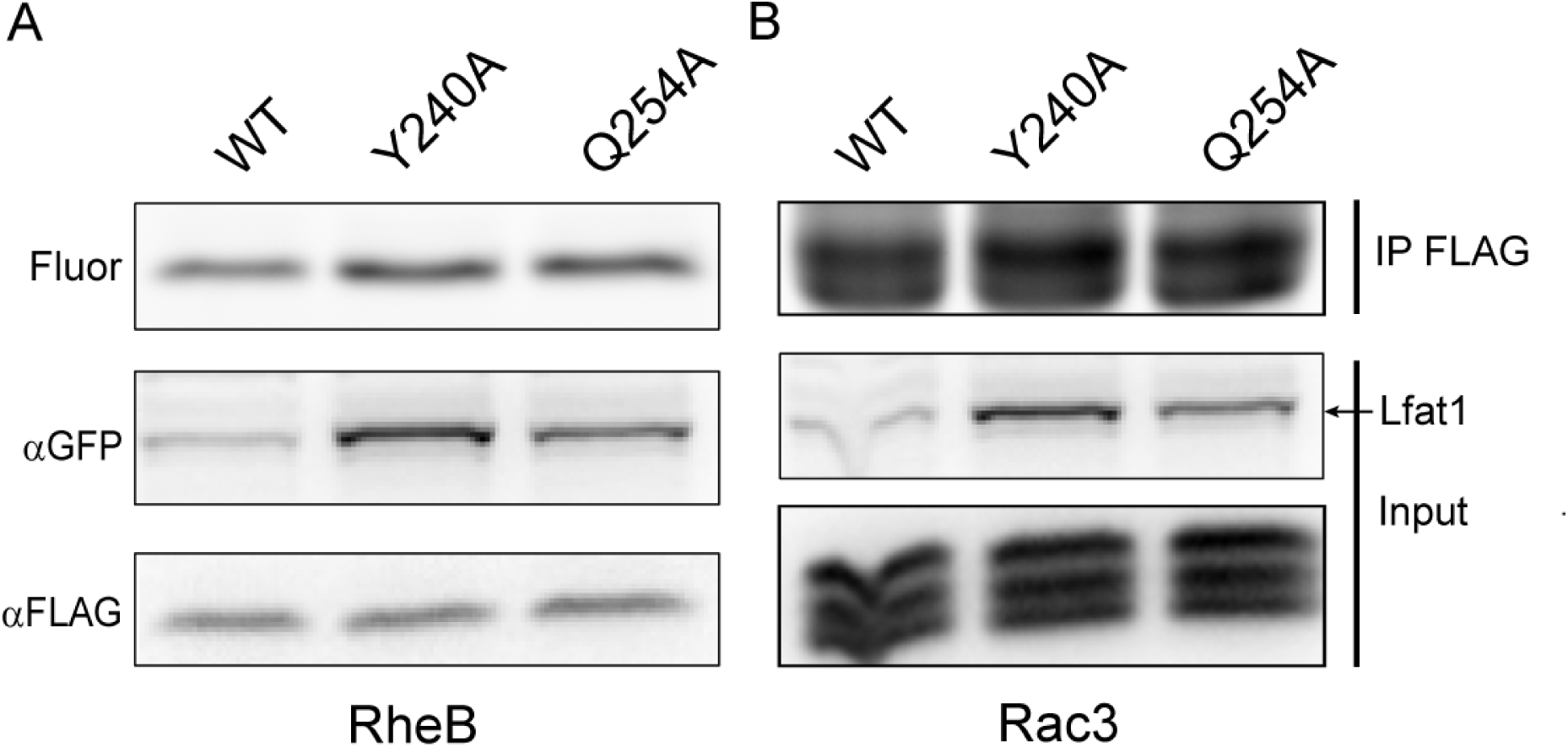
Figure supplement 2. The lysine fatty acyltransferase activity of Lfat1 does not dependent on actin binding. Lfat1 catalyzes lysine fatty acylation of RheB (A) and Rac3 (B). N-terminal Flag-tagged RheB and Rac3 were co-expressed with either GFP-Lfat1 WT, Y240A, or Q254A mutant in HEK293T cells for 24 hours. Flag-RheB and Flag-Rac3 were enriched using immunoprecipitation and subjected to Click chemistry. The samples were then separated on SDS-PAGE and scanned for fluorescence signals or analyzed by Western blot.

## Notes

### Competing Interest Statement

The authors have declared no competing interest.

### Summary of Updates

Figures 3E was modified with quadratic fitting.

